# Accelerating Cell Culture Media Development Using Bayesian Optimization-Based Iterative Experimental Design

**DOI:** 10.1101/2024.10.29.620971

**Authors:** Harini Narayanan, Joshua Hinckley, Rachel Barry, Brendan Dang, Lenna A. Wolffe, Adel Atari, Yuen-Yi (Moony) Tseng, J. Christopher Love

## Abstract

Optimizing operational conditions for complex biological systems used in life sciences research and biotechnology is an arduous task. Here, we have applied a Bayesian Optimization-based iterative framework for experimental design to accelerate cell culture media development for two applications. First, we show this approach yields new compositions of media with cytokine supplementation to maintain the viability and distribution of PBMCs in culture. Second, we applied this framework to optimize the production of three recombinant proteins in *K.phaffii* cultivations. For both applications, we identified conditions with improved outcomes compared to the initial standard media using 3 to 30 times fewer experiments than other methods such as the Design of Experiments. Subsequently, we also demonstrated the extensibility of our approach to efficiently account for additional design factors through transfer learning. These examples demonstrate how coupling data collection, modeling, and optimization in this iterative paradigm, while using an exploration-exploitation tradeoff in each iteration, can reduce the time and resources for these types of optimizations.

## 1. Introduction

Cell culture is an essential technique used throughout life sciences to study cellular, molecular, and disease biology. It is also a critical unit operation in biotechnology used to manufacture a wide range of products such as therapeutics, food proteins, peptides, biofuels, metabolites, industrial enzymes, biomass, and cells themselves for cell therapy, and the artificial meat industry^1,2^. The medium provides the essential nutrients and elements required for the growth and proliferation of the cells, the production of intended compounds, and the quality of the product^3,4^.

Optimizing the compositions of these media is a common challenge across applications. The components required are manifold, including nutrients, amino acids, nitrogen sources, carbon sources, salts, and growth hormones, among many others. This diversity presents a highly combinatorial design space with complex interactions^2^. Additionally, selecting the most suitable media significantly depends on the type and lineage of the cells, the objective of the cell culture (homeostasis, growth, differentiation), and the required operating conditions. These factors together render this task a resource-intensive and laborious one. To address this optimization, standard practices across fields rely on either the use of well-documented, historical formulations of ‘universal’ standard media^3^ or formulations resulting from a limited optimization using one factor at a time (OFAT), or a statistical Design of Experiments (DoEs)^5–9^.

Algorithms for metaheuristic optimization have been used as an alternative to these conventional approaches^3,10^, particularly for cases of bacterial cultivation and fermentation applications^2,3,11^. These algorithms are often combined with surrogate models, such as quadratic response surface methodology (RSM), Artificial Neural Networks (ANNs), or tree-based approaches^12^, to represent the underlying relationship between design factors and the target objective. These studies, however, decouple data collection, modeling, and optimization. Data are first collected using one of the statistical DoE approaches, and subsequently used for model development followed by optimization focused on maximizing or minimizing a desired target^2,3,12^. This staging requires significant coverage of the design space with the collected data to build robust surrogate models and avoid local regions of suboptimal solutions. Furthermore, these approaches have a limited ability to representatively account for the intrinsic noise in biological datasets when developing the surrogate model and performing the optimization^3^. Finally, all the methodologies, including OFAT and DoE, use only two types of design factors (continuous and discrete)^3,13^. Certain media components present multiple formats from which to choose, such as the type or source of carbon (e.g., glucose, glycerol, lactate, fructose, etc.) or nitrogen (e.g., ammonium salts, urea, glutamine). This representation of identity introduces categorical design factors that OFAT and DoE are not designed to accommodate^3,13^.

To address these challenges, we demonstrate here an iterative approach to experimental design that relies on Bayesian Optimization (BO)^14^. This strategy provides two key benefits. First, the use of a probabilistic surrogate model (Gaussian Process (GPs)^15^) is particularly well-suited for biological applications. This type of model can include prior beliefs about the system, incorporate process noise in its implementation, and obtain confidence in its predictions by associating higher uncertainty with unexplored parts of the design space. Additionally, GP models allow alternative encoding for categorical variables other than one-hot encoding approaches (OHE), which is generally regarded as an inefficient formulation as it increases the dimension of the data and adds sparsity, making the model training inefficient and increasing data requirement. Second, while planning new experiments, BO can encode a trade-off between probing unexplored regions of the design space (Exploration) and refining previously identified regions favored for the target objective(s) (Exploitation). This balance between exploration and exploitation ensures scouting of the unexplored design space, inherently minimizing the impact of local optima. This feature also dictates the planning of experiments to meet a certain objective, avoiding extensive characterization of unfavorable regions. As a result, the overall experimental burden can be reduced, accelerating the optimization. For these reasons, BO has been applied to various applications beyond computer science and robotics such as protein engineering^16–18^, reaction optimization^19^, synthetic gene design^20–22^, material science^23,24^, drug formulation^25^, and process optimization^26,27^.

We demonstrate the application of a BO-based framework to efficiently optimize the composition of cell culture media for two distinct use cases relevant to life sciences and biomanufacturing. In one case study, we show the optimization of a media composition that maximizes the viability and maintains the phenotypic distribution of peripheral blood mononuclear cells (PBMCs) *ex vivo* for up to 72 hours. In a second example, we applied this approach to determine a medium to maximize recombinant protein production by the yeast *Komagataella phaffii* (*K. phaffii*). For both applications, improved performance was achieved compared to current standard media conditions with up to 3-fold reduced experimental burden compared to the state-of-the-art DoE approaches. The reduction of experimental burden was further magnified with the increasing number of factors resulting in a 10- to 30-fold reduction when considering 9 design factors. Additionally, such a framework facilitates the transfer of learning and allows for modifications to the design space such as adding new media supplements. These results show how a BO-based active learning approach to media optimization could improve performance in cell culture for specific objectives and support additional mechanistic studies on key factors and interactions within these systems.

## 2. Results

The workflow for the BO-based active learning involves both experimental feedback and model training that reinforces the prediction of the target objective (Fig. 1A). The algorithm starts by planning and performing an initial set of experiments to build the first implementation of the surrogate GP model. The GP subsequently interacts with the Bayesian optimizer, which informs the next set of experiments that are designed to balance both exploration and exploitation of the design space. With each new dataset, the GP model is updated, and the process continues until the model converges (or the experimental budget is spent). The studies here focus on optimizing a biological objective (e.g., cell viability, titers) as a function of the composition of media.

**Figure 1:**
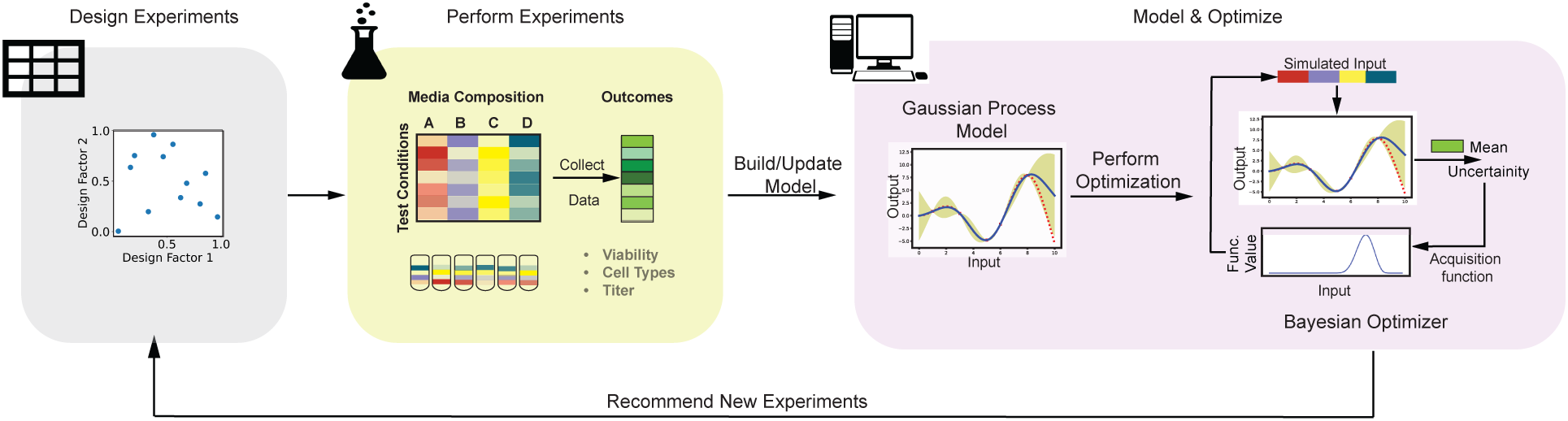
Workflow. (A) Schematic representation of the BO-based iterative experimental design workflow.

### 2.1. Optimization of media for homeostatic culture of PBMCs ex vivo

PBMCs are critical raw materials and yield data used for drug development, disease monitoring, and therapeutics. Examples include studying drug cytotoxicity^28^, co-culturing with solid tumors to understand the role of the tumor microenvironment^29,30^, and applications focusing on differentiated subpopulations of the immune cells such as T cells^31^ or NK cells^32^ for immunotherapies^33^. It is, however, difficult to maintain these primary cells in culture for extended durations with typical commercial media without losses in viability or changes in the distribution of cell types. We sought to apply our BO-based approach to perform two sequential optimizations. First, we aimed to determine a media blend of commercially available media (DMEM, AR5, XVIVO, and RPMI-10) that would maximize cell viability. Second, we undertook an optimization using cytokines and chemokines to achieve a balance of key lymphocytic populations representative of the *ex vivo* distributions (Fig. 2A).

**Figure 2:**
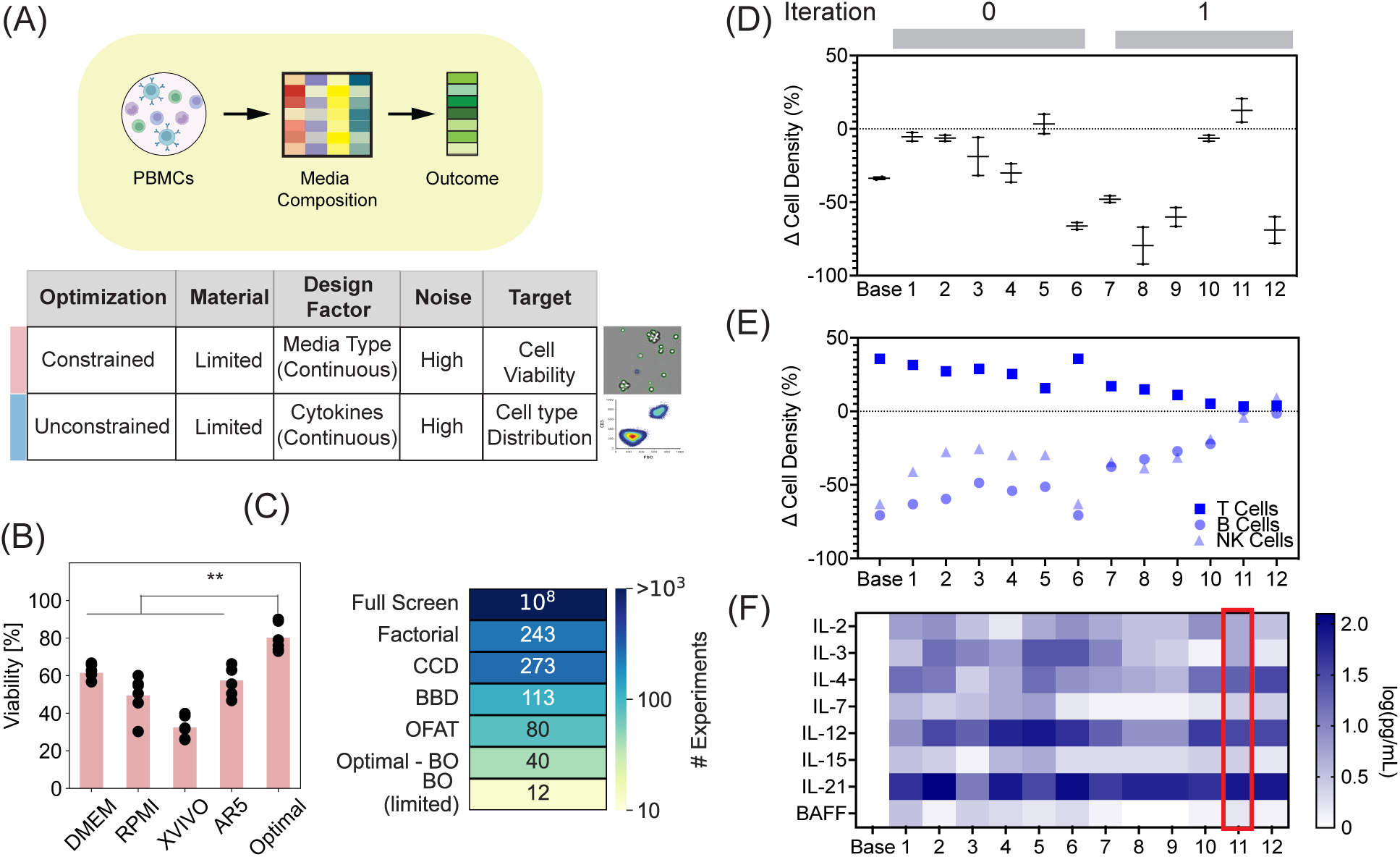
Media blend and cytokine composition optimization PBMC cultures. (A) Workflow and study parameters to optimize composition of media to maximize viability and maintain homeostasis of PBMCs in cultures (B) PBMC cell viability as a function of the optimized media blend compared to individual standard media (DMEM, RPMI, XVIVO, and AR5). (C) Comparison of the number of experiments to execute the different strategies for designing experiments considering 8 different cytokines (D) Change in total cell density before and after 3 days of PBMC culturing using different compositions of cytokines with the red arrow indicating the condition meeting the desired objective (E) Change in cell density of subpopulations of lymphocytes before and after 3 days in culture under different cytokine compositions (red arrow indicates the condition meeting the desired objective) (F) Composition of the different cytokine compositions tested (red box corresponds to the condition meeting the desired objective)

We hypothesized that different commercial formulations of media have different sets and (or) quantities of nutrients, hormones, and growth factors, and combining these in different ratios could yield a new composition capable of maintaining high viabilities (>70%). This framing yields a constrained optimization problem with continuous design factors, such that the contribution of the different media in the blend sums to 100%. We applied our BO-based iterative design approach to maximize the cell viability of healthy PBMCs after 72 hours in culture. With 24 experiments, an optimized blend of the media showed a statistically improved viability (75-80%) compared to individual media (Fig. 2B).

Unsurprisingly, we found that the blend of commercial media yielding improved viability of PBMCs did not uniformly maintain the diverse subpopulations of lymphocytes, favoring T cells over populations of B cells and NK cells. To address this imbalance, we selected eight design factors to test based on their known roles in homeostasis, including interleukins (IL-2, IL-3, IL-4, IL-7, IL-12, IL-15, IL-21) and B-cell activating factor (BAFF)^34,35^. The scope of this design space is substantial and would require large numbers of experiments to evaluate using DoE or OFAT (Fig. 2C). This trait, therefore, makes performing such optimizations on sparse clinical samples challenging for many applications like drug evaluations in precision medicine or the production of non-T-cell-based cell therapies. Using our approach, we sought to identify combinations of the additional media supplements that would maintain the distributions of cell types after 72 hours of cultivation compared to the *ex vivo* distribution. With as few as 12 additional experiments, we found a combination that retained both the cell density (numbers) and the distributions of cell types (Fig. 2D, E, F). Interestingly, similar mixtures of the supplements (Expts. 11 and 12), differing predominantly in the concentration of IL-3, showed a large difference in total cell densities despite similar distributions of cell types as observed *ex vivo*. The optimized combinations also showed reduced concentrations of two cytokines commonly used in media for CAR-T cell cultures (IL-7 and IL-15) compared to the initial designs tested (Expts. 1-4). These modest differences observed in the total composition highlight the nuances of formulating media for primary cells.

### 2.2. Enhancing recombinant protein production by *K. phaffii* with carbon source supplements

Optimization of media is also important for applications in industrial biotechnology. As a second example, we applied our BO-based active learning approach to improve recombinant protein production by a yeast host^36^, *K.phaffii*, commonly used to produce food proteins, materials, and biologics. Standard compositions of media used for this cultivation rely on either a complex^37^ or basal salt media^38^ supplemented with glycerol^39^ or glucose^40^ for biomass accumulation, followed by methanol with or without sorbitol co-feeding to induce production. These formulations (and their use in fermentation) follow canonical standard protocols^41^. We hypothesized that optimizing the concentrations of carbon sources in small-scale cultures could improve the production of secreted proteins. We devised a design space including four design factors: the concentration of glycerol during biomass accumulation, the concentration of methanol during production, the type of carbon co-feed supplement in production (categorical factor), and the concentration of the co-feed. To evaluate the generalizability of our approach and compare convergent solutions, we optimized the required carbon sources for three different proteins with varied complexities: an engineered variant of the SARS-CoV-2 RBD subunit (RBDJ)^42^, Human Serum Albumin (HSA), and an IgG1 monoclonal antibody trastuzumab (Fig. 3A). We selected two commonly used carbon conditions as references to compare the resulting compositions (Benchmark1 and Benchmark2 in Table 1).

**Figure 3:**
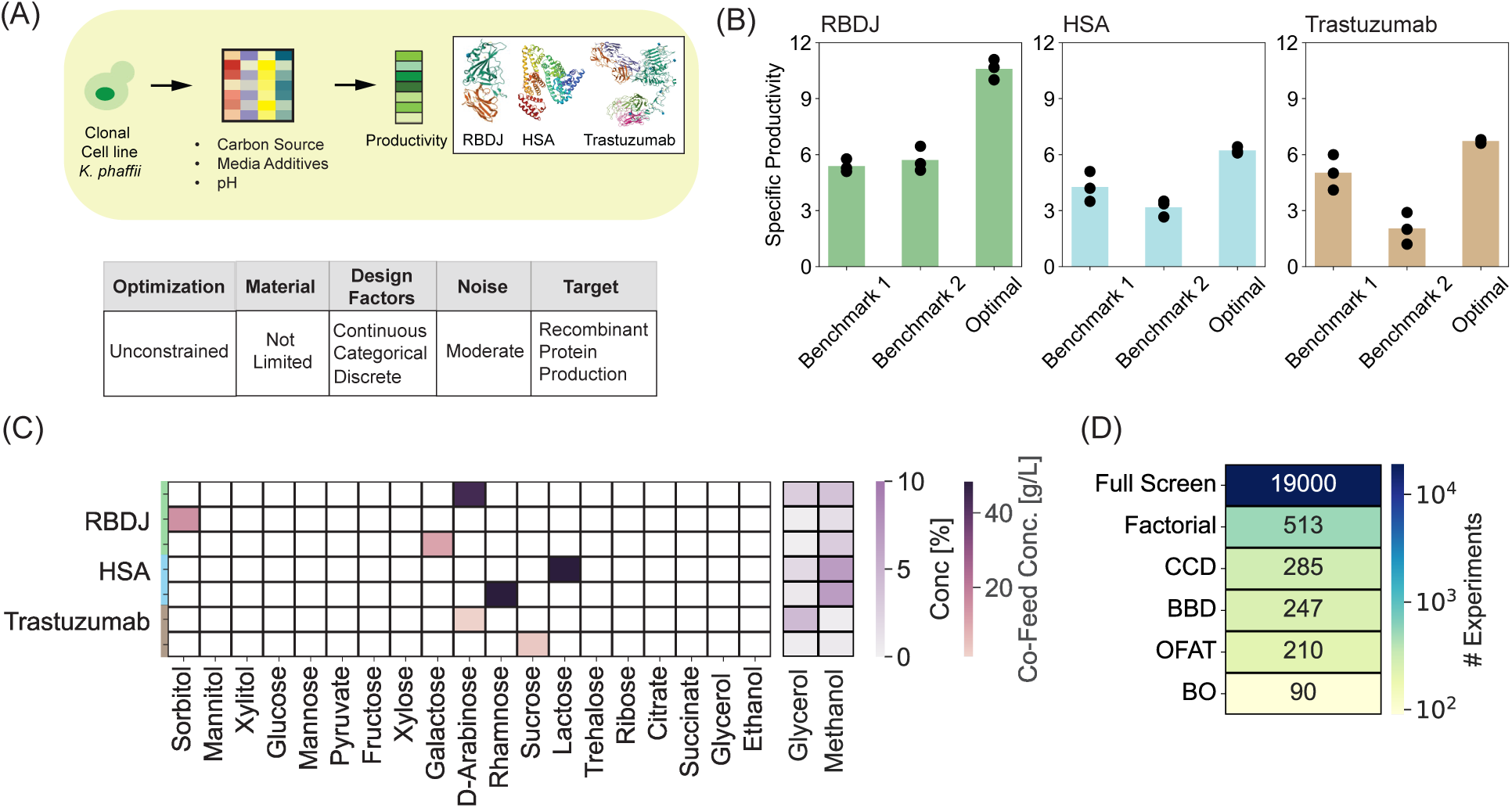
Carbon source optimization for recombinant protein production in *K.phaffii*. (A) Workflow and study parameters for optimizing media supplements to maximize recombinant protein production in cultures of *K.phaffii*. (B) Comparison of specific productivity using optimized media compositions and 2 benchmark conditions for RBDJ, HSA, and trastuzumab. (C) Optimized compositions of carbon sources and specific productivities for the respective proteins. (D) Comparison of the number of experiments to execute the different strategies for designing experiments.

**Table 1:**
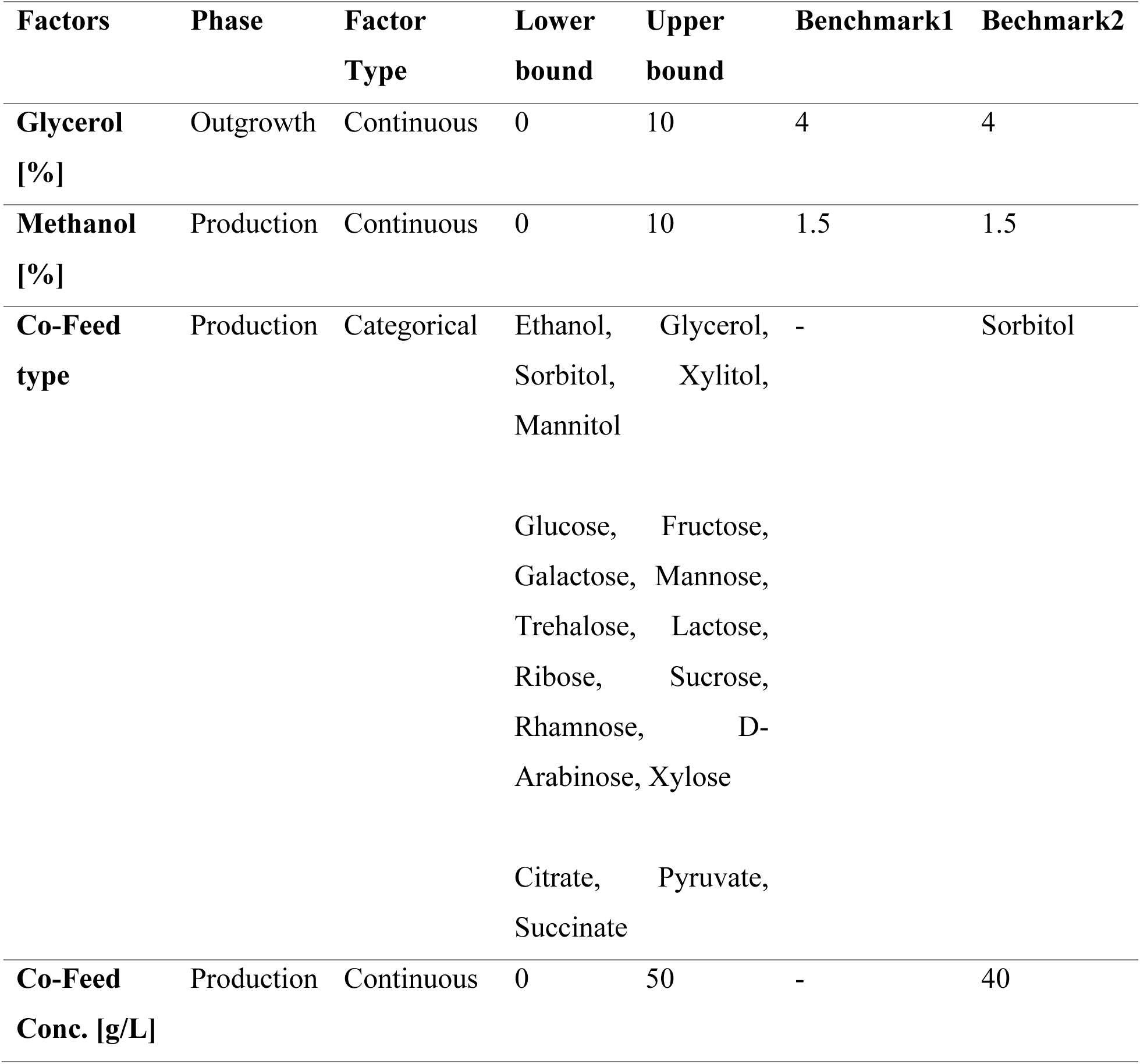
Factors considered in the experimental design and the corresponding ranges.

To account for the categorical variables in our surrogate GP model, we adopted a modified kernel design instead of using OHE. To validate the choice, we retrospectively compared the predictive accuracy of the GP models trained with our designed kernel and OHE (Fig. S1A). We confirmed that the average errors made by GP with the designed kernel were 33-50% smaller compared to OHE. We assessed convergence here as the agreement between the model prediction and the experimental observation (Supplementary information).

We then demonstrated that the BO could identify optimized combinations of carbon sources to improve the specific productivity (measured by mg/L/OD600) relative to the benchmarks, albeit there were different degrees of improvement for each protein (Fig. 3B). For each molecule, we identified multiple and distinct compositions of media (3 compositions for RBDJ, 2 for HSA, and 2 for trastuzumab) that could optimize protein production to a similar degree (Fig. 3C). These media yielded a 2.5-fold improvement in specific productivity compared to both the Benchmark media for RBDJ (12 mg/L/OD600 vs 5 mg/L/OD600). For HSA, we observed a 2-fold improvement compared to Benchmark2 (6 mg/L/OD600 vs 3 mg/L/OD600) but only a 1.5-fold improvement compared to Benchmark1 (6 mg/L/OD600 vs 4 mg/L/OD600). Finally, for trastuzumab, we observed a 3-fold improvement compared to Benchmark2 (6 mg/L/OD600 vs 2 mg/L/OD600) and no statistically significant improvement compared to Benchmark1. These differences in observed improvements suggest there are unique protein-dependent bottlenecks faced by the host that cannot be alleviated by only optimizing carbon sources.

Furthermore, the optimal media compositions differed considerably among the tested molecules with no two molecules converging to the same composition (Fig. 3C), highlighting the unique requirements faced when optimizing media for the efficient production of different recombinant proteins. Given this trait, it is ideal to develop new protein-specific media without requiring excessive resources or time. Using our BO-based active learning approach, we found we could optimize the carbon sources required using only 90 experiments over 7 experimental iterations, requiring a total of ∼1 – 1.5 months (Fig. 5A). This total experimental load was ∼ 2.5 – 3 times lower than the predicted number of experiments required for standard designs of DoE and several orders of magnitude lower than a comprehensive search to screen the design space (Fig. 3D).

### 2.3. Elucidation of the algorithm characteristics: Exploration-Exploitation tradeoff

Having demonstrated the capabilities of the BO algorithm for accelerated, resource-efficient media optimization, we next sought to elucidate the characteristics of the algorithm by investigating how the algorithm navigates the selected design space and its corresponding impact on the target. First, the use of a space-filling design to generate the initial iteration of experiments maximized the variability in the input-output combinations seen by the GP model, thus, allowing for an efficient initial representation of the system. The wide coverage of the design space tested in the initial iteration for the PBMCs (Fig. 4B) is reflected in the range of cell viabilities measured (ranging from 5 % to 62%) (Fig. 4A). The diversity of the initial assessment is also explicitly evident from the varied ratios of the different media types (DMEM, AR5, XVIVO, and RPMI-10) in the designed initial blends (Fig. 4C). We note that the constraint on media blending (that is, the sum of the ratios of different media type should equal 1) in this example limited the accessible region to design experiments (Black dashed line, Fig. 4B). Similarly, the initial space-filling design to optimize the carbon sources for the yeast cultivations was also broadly distributed (Fig. 5B) subsequently resulting in a wide distribution of specific productivities measured (ranging from 0.1 to 9 mg/L/OD600) (Fig. 5A).

**Figure 4:**
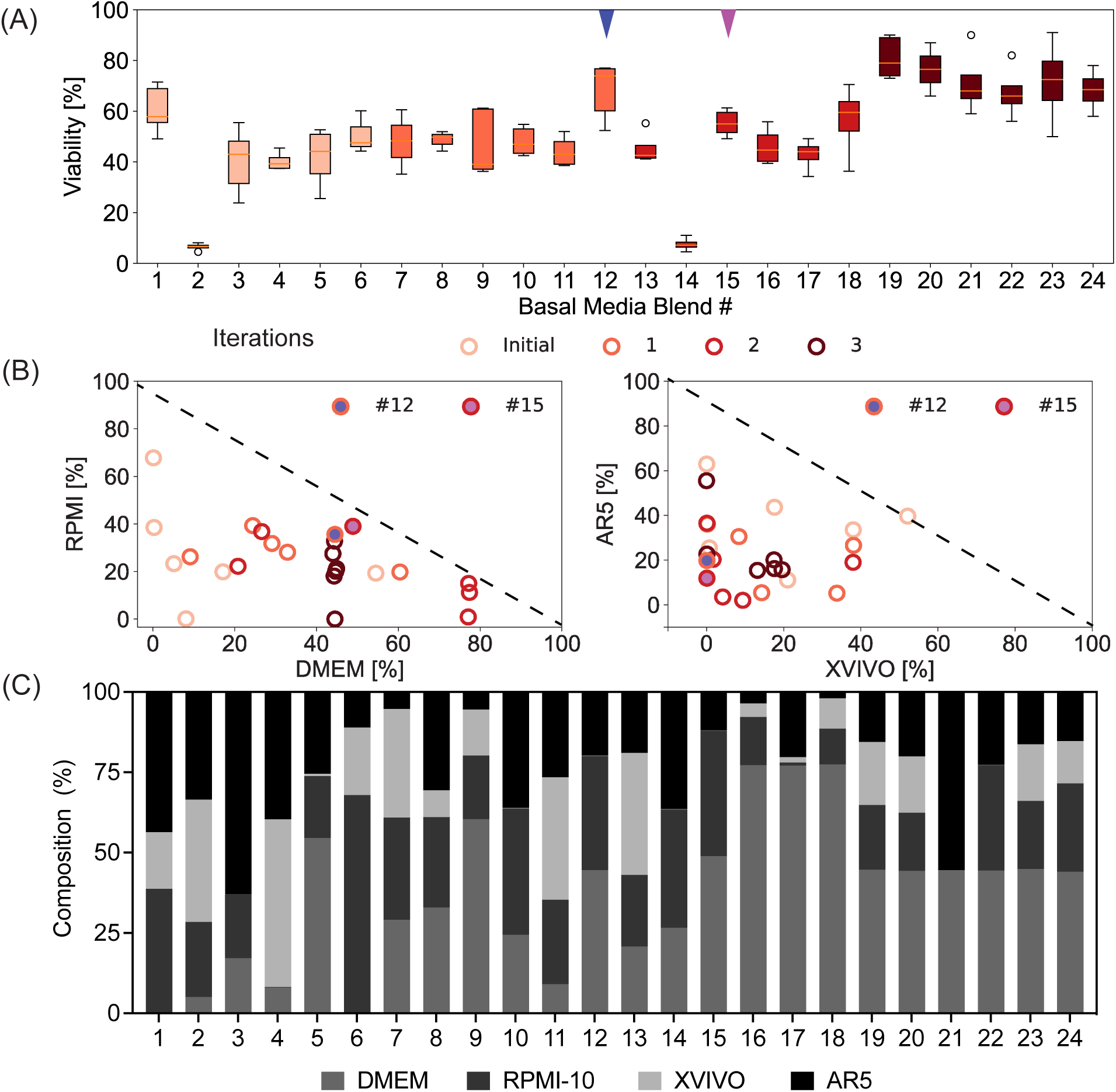
Characterization of exploration-exploitation trade-off for PBMC media blending case. (A) Evolution of cell viability of the experiments planned in the different iterations. (B) Evolution of the location of the experiments in the feasible design space during the different iterations. (C) Composition of the different media blends tested in the different iterations.

**Figure 5:**
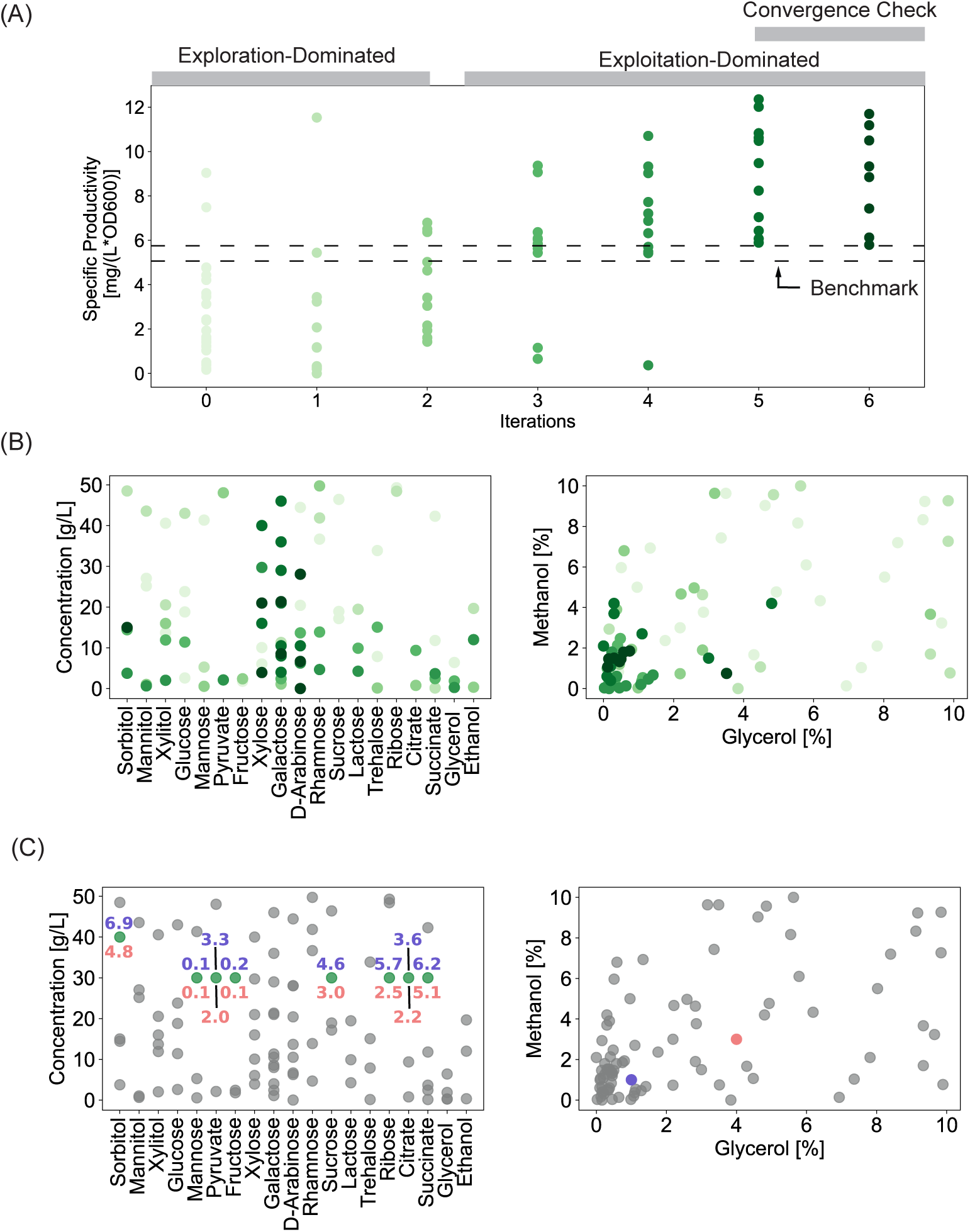
Exploration-exploitation trade-off and confirmation of poor specific productivity in the region lacking experiments. (A) Evolution of specific productivity distribution of the experiments planned in the different iterations. (B) Evolution of the location of the experiments in the design space in the different iterations. (C) The carbon source type and concentration of the validation experiments planned (Green Circles). The numbers represent the specific productivity with the colors indicating the corresponding glycerol–methanol conditions in D (D) Scatterplot of the percentages of methanol and glycerol tested. Pink and purple circles – two combinations of glycerol and methanol tested with the selected carbon source type and concentrations marked in Fig. 5C.

In each future iteration, the optimizer attempted to find a trade-off between planning experiments in unexplored regions of the design space and refining its confidence in regions identified as favorable for the target objective (exploitation). Practically, given that the unexplored regions represent a larger portion of the design space at the start of the sequential campaign, we confirmed that the recommended experiments planned in early iterations of the optimization favored exploration, and then progressively moved towards exploitation-dominated designs.

For the example of media blending for PBMCs, we observed that exploration dominated the first two iterations (Iterations 0 and 1), with cell viability varying from 5% to 75% over wide coverage of the feasible design space. Subsequently, Iteration 2 included a mix of exploration and exploitation: For instance, Blend 15 exploited a previously observed region covered by Blend 12. Subsequently, the final iteration (Iteration 3) exploitatively reduced the search space to a specific ratio of DMEM with a focus on perturbations involving different combinations of the other media types (Fig. 4B, C).

For the optimization of media for protein production with yeast, we similarly observed a large distribution of values of specific productivity (Fig. 5A) with widespread spacing of experiments covering the design space in Iteration 1 and 2 (Fig. 5B). Unsurprisingly, the algorithm planned experiments in Iteration 1 for types of carbon sources not probed in the initial experiments since these regions of the design space still had the highest uncertainty, dominating the objective function. The subsequent iterations (3-6), however, emphasized exploitation as observed by the higher fraction of experiments planned in a limited region of the design space resulting in increased specific productivity (Fig. 5A) with limited testing of other regions of the design space (Fig. 5B).

These analyses show the algorithm progressively learned the favorable/unfavorable regions of the design space as it reduced the experimental testing in the unfavorable regions. For the PBMC culturing, this progression resulted in a few experiments being planned for media blends with XVIVO > 60%. Reviewing the tested media blends, it is clear that blends dominated by higher XVIVO amounts (Blends 2, 4, 11, and 13 – Fig 4C) resulted in lower cell viabilities with the best viability of only 40% (Fig. 4A).

Similarly, for the production of RBDJ by *K.phaffii*, specific carbon source types (e.g., mannose, pyruvate, ribose, glycerol, etc.; especially in the concentration range of 20-40 g/L) were restrictively probed (Fig 5C). To validate that these ignored parts of the design space would result in poor specific productivity, we manually picked 16 different conditions with the eight different carbon sources from these regions of the design space and tested the specific productivity of RBDJ under those conditions (Fig. 5C). The eight carbon sources were tested with two different glycerol and methanol concentrations (Fig. 5D). The specific productivities for all these selected conditions ranged from 0.1 to 6 mg/L/OD600 and were lower compared to the algorithm-identified optimal conditions for RBDJ (specific productivity of 12 mg/L/OD600) (Fig. 3B).

Taken together, these analyses on the iterative progression of the models for both cases tested support the ability of BO to accelerate optimization and minimize resources used. The synergy between experimentation, model building, and optimization in this iterative framework, coupled with the gradual trade-off between exploration and exploitation in each iteration reduces the number of experiments allocated to less optimal regions of the design space.

### 2.4. Extending the approach with Transfer Learning to incorporate additional media supplements

Both cases tested here yielded improved performance for the respective objectives subject to the selected media components. It is apparent in both cases and, more generally, that performance could be further enhanced by considering additional influential factors in the optimization. It is, therefore, desirable to allow continued improvement and extensions of the model to incorporate new design factors or objectives, while using the learnings generated from the existing data to minimize future work to extend the optimization. This goal requires capabilities to transfer learnings to modified design spaces (e.g., expanding the range of the design factors or adding additional factors) or to alternative biological systems (e.g., production of other molecule types or culturing blood cancer cell lines). In this work, we explored the feasibility of extending our current framework to the first case. GP models are well-suited to this extension since they can learn from data through the context of similar experiments in the design space. We sought to demonstrate this feature by considering five additional media additives to optimize the recombinant protein production in *K.phaffii* (Table 2), using HSA as a test protein because it showed only moderate improvement when only the selection of carbon sources was considered.

**Table 2:** Additional factors considered in the experimental design of the transfer learning study and the corresponding ranges.

| **Factors** | **Phase** | **Factor Type** | **Lower bound** | **Upper bound** | **Benchmark** |
| --- | --- | --- | --- | --- | --- |
| **Glutathione [mM]** | Outgrowth, Production | Continuous | 0 | 50 | 0 |
| **pH [-]** | Outgrowth/ Production | Discrete | 5.75, 6.0, 6.5 |  | 6.5 |
| **Tween20 [%]** | Outgrowth, Production | Continuous | 0 | 10 | 0 |

To seed this new iteration of the model that included the additional supplement, we used the current GP as the prior (instead of using a space-filling design) and used the optimizer to determine subsequent experiments. We included all nine factors (the new supplements and the four prior ones) in designing new experiments, allowing the model to re-learn dependencies in the modified design space as needed. Iterating in this way yielded a modified composition of media that improved the specific productivity for secreted HSA from 6 mg/L/OD600 (starting point) to 13 mg/L/OD600 (Fig. 6A). This realized improvement corroborates our hypothesis that hosts producing recombinant heterologous proteins may require specific tailored media composition to maximize their specific productivity due to unique metabolic requirements or protein-dependent features (folding, assembly).

**Figure 6:**
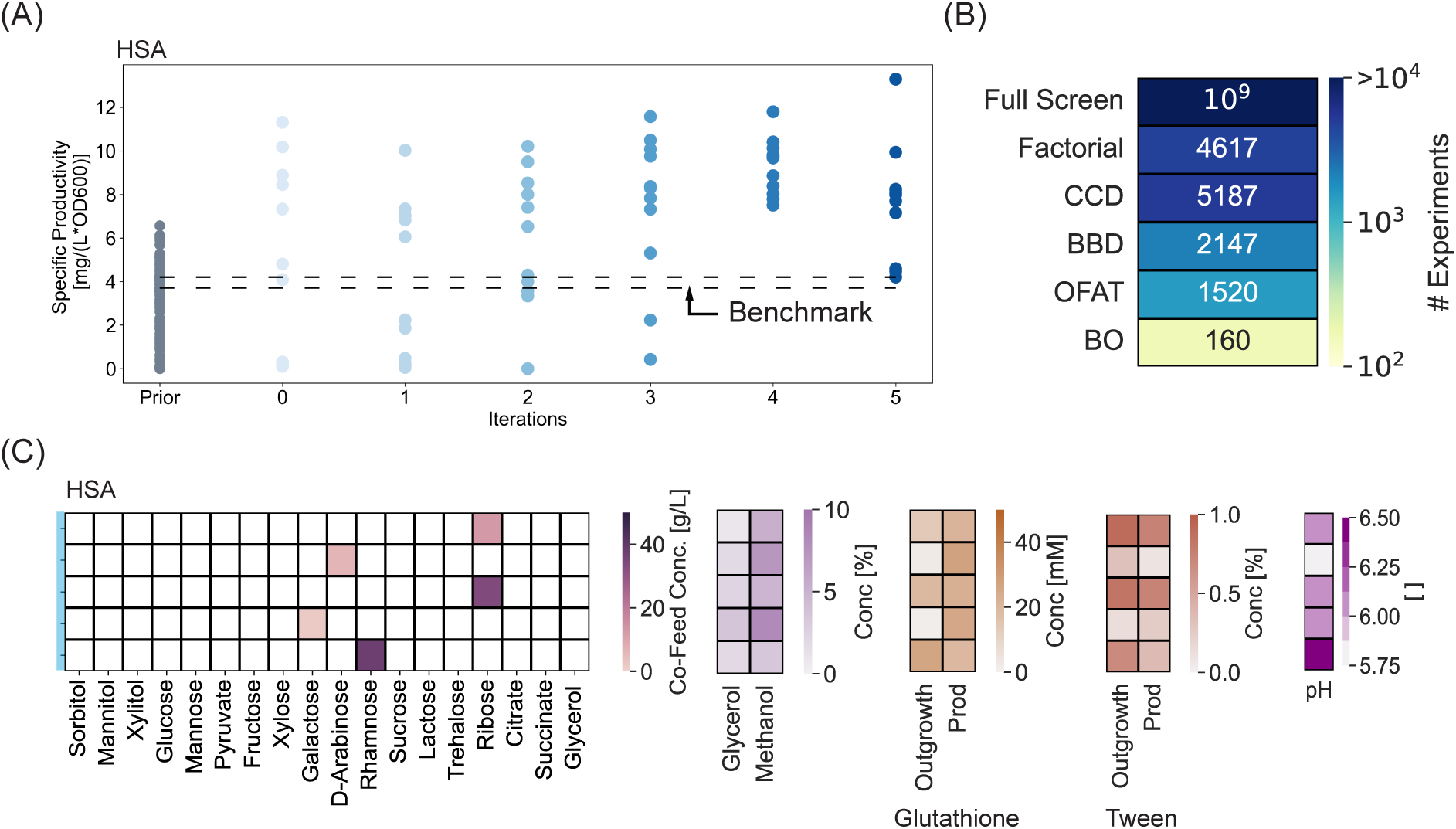
Transfer learning to a new design space with additional factors. (A) Evolution of specific productivity distribution of the experiments planned in the different iterations. (B) Comparison of the number of experiments to execute the different strategies for designing experiments. (C) Composition of media supplements for top 5 candidates with highest specific productivities in decreasing order.

One impact of starting the model with the GP from the prior task is that the algorithm minimized the experiments planned for certain carbon sources that had yielded poor specific productivities previously (e.g., sorbitol, mannitol, xylitol, glucose, mannose, succinate, and glycerol), and focused on a subset of alternative carbon sources within the first two rounds (Fig. S2). Similarly, starting from the initial iteration, the algorithm planned most of the experiments with concentrations of glycerol < 5% and those of methanol between 1.5 – 8%. As a result, new optimized media conditions required only 72 additional experiments (including 12 to confirm model convergence) to consider the new design space of 9 factors. The two tasks combined required a combined total of ∼160 experiments—several orders of magnitude lower than a practically infeasible full screen and ∼ 10- to 30-times fewer experiments than simple OFAT and traditional DoE approaches, respectively (Fig. 6B).

The experimental burden of DoEs increases substantially as the design space expands, and consequently, often only a subset of factors are considered in any given optimization to reduce the overall number of experiments performed^10^. For instance, in our nine-factor study here, a study might hold the prior optimized conditions for the carbon source constant, and simply perform a separate DoE for the five additional design factors. Interestingly, in the case we considered, however, this common approach would not have yielded the best productivity: The new media supplements resulted in an alternative preference of carbon sources for maximizing the productivity (from Rhamnose and Lactose to Ribose, D-Arabinose, and Galactose) (Fig. 3C, 6C). Whereas the use of Rhamnose as a co-feed resulted in high specific productivity (∼10.5 mg/L/OD600), the concentration was altered from that found with the four-factor design space. These data support the flexibility of this BO-based active learning approach for media optimization to accommodate the posterior addition of design factors (such as new additives) or expand the concentration range of existing design factors^25^. This ‘bootstrapping-type’ approach to optimization would become increasingly valuable as the design space expands, as is often the case in media optimization when considering several types and concentrations of additives.

## 3. Discussion

In this study, we have presented an accelerated and resource-efficient approach for media development using Bayesian optimization-guided iterative experimental design. Using two unique experimental systems, we have shown its capabilities for cell cultures used in common applications in both life sciences and biomanufacturing. First, we optimized the media composition to maximize viability and maintain homeostasis of PBMCs in culture. In the second case, we optimized the concentrations of additional supplements used in cultivation media for *K.phaffii* to maximize the production of three different recombinant proteins. In both cases, improved performance was achieved compared to current standard media conditions with at least a 3- (e.g., 4 design factors) to 30-fold (e.g., 8-9 design factors) reduced experimental burden compared to the state-of-the-art DoE approaches. Inferring from the selection of experiments within the design space during iterative rounds, it is evident that this efficiency results from the capability of the algorithm to focus the experimental effort on identifying favorable regions for the targeted objective via a tradeoff between exploration and exploitation, and therefore, minimizing the experimental efforts in undesirable regions, reducing the overall time and resources required.

The examples of optimizations performed here, including the incorporation of transfer learning to extend the design space, show the potential for this BO-based active learning strategy to minimize experimental costs and time for complex biological tasks like media optimization. The extensibility of the models makes it possible to add new design factors or objectives without introducing artificial constraints or biases needed to manage the budget for experimental exploration of the large design space. GPs used for the models intrinsically provide an efficient way to accommodate additional media supplements or objectives a-posteriori since they associate unexplored regions of the design space (the influence of new added factors/goals) with a larger uncertainty. This approach should provide benefits for adapting models to new systems or tasks where the input materials available for experiments impose a natural limitation on how many experiments are feasible during optimizations (e.g., development of primary cancer cell culture from tissue biopsies, pediatric cell therapies).

This approach derives its advantage from coupling data collection (design), modeling, and optimization into a comprehensive, iterative process, thus strategically navigating the design space based on experimental feedback. In contrast, DoE approaches offer static designs irrespective of design factors or target types, with no active feedback based on data collected from the experimentation. Additionally, our approach offers more generalized capabilities that can account for different types of design factors and optimization tasks compared to DoEs, which can only work with discrete or continuous and cannot generate designs for constrained spaces. Finally, the approach here offers broader coverage of a design space compared to DoE studies that only test the corner and center points of the design space. The case studies presented here together invoked a range of scenarios that account for different types of design factors (continuous, discrete, categorical), frameworks for optimization (constrained, unconstrained), degrees of biological noise (moderate to high), and intrinsic limitations on the material (cost, time). One disadvantage of our current implementation is the procedure for *in silico* sampling needed to accommodate categorical variables relies on randomly sampling many potential experimental suggestions, increasing computational time (∼ 2-4 mins/experiment). Using recent advancements in optimization approaches^43^, the efficiency of this sampling process could be improved.

In conclusion, we have focused here on a BO-based active learning approach to media optimization and demonstrated improved performance in cell culture for specific objectives. These examples, however, highlight the potential for this approach to extend to other complex biological applications such as process development. The extensibility of the strategy suggests that incorporating such BO-based experimental design as a standard practice in life sciences research could facilitate both the generation of foundational data to support new predictive models in biological systems and further accelerate development pipelines for new systems. For example, the models developed here to optimize media for PBMCs could be extended to address perpetual cell lines and adapted for patient samples in cases where the materials are limited (e.g., pediatric cancers). Similarly, media and process development for new protein molecules or engineered strains could be accelerated by starting from the models and data generated here. Related applications like scaling up process conditions across systems could also benefit from this strategy for optimization which uses synergy between iterative experimental data and modeling to inform optimization. We postulate that this approach and similar ones like reinforcement learning could establish modular frameworks for improving predictive capabilities across multiple complex biological systems.

## 4. Methods

### 4.1. Experimental Methodology

#### 4.1.1. Culturing PBMCs

Peripheral blood mononuclear cells (PBMCs) were thawed in a 37°C water bath, followed by adding the thawed cells to 4 mL of DMEM supplemented with 10% fetal bovine serum (FBS). The cells were centrifuged at 500 x g for 5 minutes, the supernatant was discarded, and the pellet was resuspended in 2 mL of fresh DMEM. Cells were counted, and the appropriate volume of cell suspension was calculated to achieve a final concentration of 1 × 10^6^ cells/mL in 2.3 mL of the chosen reactor medium (determined based on BO suggested experimental design). After centrifugation at 500 x g for an additional 5 minutes to remove any residual DMEM, the cells were resuspended in 2.3 mL of the reactor medium. A volume of 200 µL of the cell suspension was then pipetted into each well of the specialized C.BIRD reactor plates, ensuring that empty wells were filled with 200 µL of PBS or water and 10 mL of PBS or water was added to the plate reservoir. The functional lid was attached to the 96-well plate, and the assembly was placed in a 37°C incubator with standard settings used for cancer cell culture, including 5% CO₂ and high humidity (∼95%) to maintain pH and prevent evaporation.

#### 4.1.2. Viability Assays

Cell viability and counting were performed before and after incubation in the bioreactors. After the culture duration (72 h), cells from the replicate conditions were pooled into a single tube, and 40 µL of EDTA was added to each sample, to help prevent aggregation and detach PBMCs from the reactor surfaces to ensure accurate cell counting. A 1:1 mixture of the cell suspension and AOPI (ViaStain) was prepared, and cell counts were performed in triplicate using the Nexcelom Cellaca plate reader.

#### 4.1.3. Flow Cytometry

Peripheral blood mononuclear cells (PBMCs) were isolated from healthy donor blood samples (STEMCELL Technologies) using density gradient centrifugation and pooled into a 1.5 mL Eppendorf tube. A 100 µL aliquot of cells was extracted for viability assessment, while the remaining volume was centrifuged at 500 x g for 8 minutes. During centrifugation, 5 µL of Fc block (Human TruStain FcX; BioLegend) was added to 95 µL of FACS buffer (phosphate-buffered saline [PBS] supplemented with 1% bovine serum albumin [BSA] and 0.1% sodium azide) per sample. After centrifugation, the supernatant was discarded, and the cells were resuspended in 100 µL of Fc block solution and incubated on ice for 20 minutes. Concurrently, a staining solution was prepared by adding 5 µL of each fluorescent-conjugated antibody (anti-CD20-FITC, anti-CD45-APC, anti-CD56-APC-AF750, Zombie Violet; BioLegend) to FACS buffer, adjusting the volumes according to the number of samples. Cells were incubated on ice in the dark for 30 minutes. Following incubation, the cells were washed twice by centrifugation at 500 x g for 5 minutes and resuspended in 200 µL of FACS buffer. Flow cytometric data acquisition was performed on a Beckman Coulter CytoFlex LX, utilizing appropriate laser configurations and voltages for each fluorophore.

#### 4.1.4. *K.phaffii* Cultivation

Twenty-four well plate scale experiments (working volume of 3 mL) were performed to test the different media formulations on Benchmark plate shakers (R.T., 600 rpm). Complex media (potassium phosphate buffer pH 6.5 (unless specified otherwise), 1.34% nitrogen base w/o amino acids, 1% yeast extract, 2% peptone) was used as the base media for the cultivation with appropriate additions of carbon sources and additives as per the algorithm. Cultivations were inoculated at 0.1 OD600 from working cell banks, outgrown for 24 h, pelleted, and resuspended in fresh production media to induce recombinant gene expression. Supernatant samples were collected after 24 h of production, filtered, and analyzed using HPLC to quantify titer. SDS-PAGE was carried out as described previously ^44^ to confirm protein bands of the right size and no product-related variants. Specific productivity was defined as relative titer normalized by cell density, measured by OD600.

### 4.2. Algorithmic Methodology

#### Initial Design

A Latin hypercube sampling for the continuous variables and a random design for the categorical variables. In Latin hypercube sampling, the designs are generated such that each hyperplane of the design space has only one point in contrast to purely random sampling from a uniform distribution that results in uneven spacing of the experiments in the design space^3,25,26^. However, since LHS considers continuous unconstrained design spaces, random sampling was used for constrained design spaces (PBMC media blending study) and categorical variables (for the *K.phaffi* recombination protein production optimization case study).

#### Gaussian Processes

Gaussian processes (GPs) are probabilistic models that learn an underlying unknown black-box function (i.e., the relationship between media additive and specific productivity) by representing them as a distribution of functions. This distribution of function is characterized by a mean *m*(*x*), and a covariance function *k*(*x*, *x*′) that is dictated based on prior beliefs about the system (Eq 1).

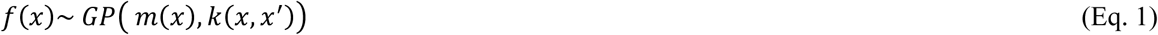

The covariance function is defined through the kernel which encodes the similarity between points in the design space and the selection of this function depends on the beliefs about the smoothness, periodicity, and trends in the design space. Since smooth trends are expected over the continuous space in this application, we used a smooth flexible Matern kernel.

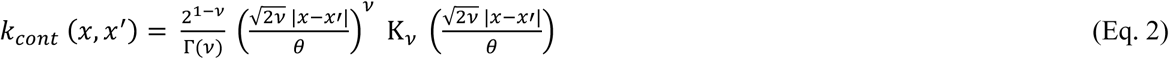

x indicates the continuous inputs, Γ is the gamma function and *κ_v_* is the modified Bessel function. The |x – x’| indicates the distance between the two points. The kernel value decreases as the distance increases beyond the length scale, *θ*. *v* is the smoothness controlling parameter and was set to 2.5 which is typical for the kernel.

For the definition of the categorical variable, we use the formulation of the categorical overlap kernel suggested by Ru et al.,^45^ that defines the kernel as the total number of categories that overlap between the two points *h_i_* and *h*′*_i_*.

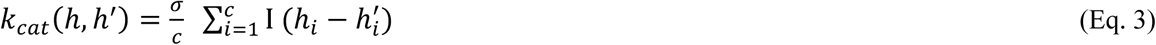

h indicates the categorical inputs, and c represents the total number of categorical variables considered (in this case, 1 - that is the type of carbon source).

The final kernel over the categorical-continuous variables together (*z*) is also adapted from Ru et al.,^45^ as follows:

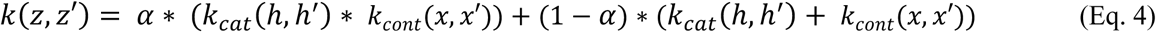

The length scale in the continuous kernel and the trade-off parameter (*⍺*) in the mixture kernel is a hyperparameter that is updated based on data to subsequently refine the priors and result in a posterior distribution.

The noise addition to function estimation (*∈_i_*) is provided through the likelihood function, in this case, Gaussian likelihood (Eq. 6).

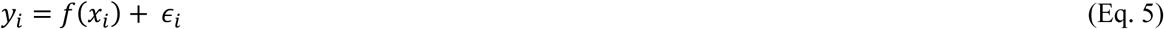

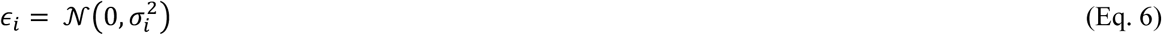

Gpy package was used to set the kernel and the Gaussian process implementation in Python.

#### Bayesian Optimization

The GP is then used by an optimizer that suggests the next set of experiments (media conditions) using an acquisition function that encodes a tradeoff between characterizing previously unexplored parts of the design space and exploiting the regions with promising targets (higher specific productivity). In this work, we use an upper confidence bound acquisition function compared to other alternatives that define the tradeoff simply using the predicted mean and uncertainty. For continuous unconstrained optimization problems (e.g., the cytokine optimization for PBMC culture) a local optimizer such as LBFGS is used to maximize/minimize the acquisition function, and a trust-region-based algorithm is used for constrained optimization (PBMC basal media blend). These optimizers are implemented through the scipy package in Python. In the presence of categorical variables (*K. phaffi* cultivation media optimization), we used a brute force approach and simulated 10000 points via LHS, and the points with the maximum/minimum acquisition function were picked.

Finally, the original implementation of BO is a truly sequential design approach planning one experiment in each iteration^14^. For most applications, as in our case, it is more practical to perform several experiments in parallel. Thus, we use a variant of BO called batch BO using the “constant liar” approach using the mean value^46^ amongst others^47,48^, which has previously shown success in other applications such as^25,26^. The batch size was determined by the throughput of the experimental system, material availability and replicate requirements.

## Authors Contributions

Harini Narayanan: Conceptualization, implementation of the algorithm, executed yeast experiments, writing manuscript

Joshua Hinckley: PBMC application design space and experimental protocol establishment, executed PBMC experiments

Rachel Barry: PBMC bioreactor cultures and flow cytometry preparations Brendan Dang: PBMC bioreactor cultures and media preparations

Lenna A. Wolffe: PBMC bioreactor cultures and media preparations

Adel Atari: PBMC bioreactor cultures and flow cytometry preparations

Yuen-Yi (Moony) Tseng: Lab facilities for PBMC work, review and revise manuscript

J. Christopher Love: Supervision, review and revise manuscript

## Acknowledgments

We thank the MIT AltHost Research Consortium for the strains of *K. phaffii* used in these studies. This work was funded with support from a SPARC grant from the Broad Institute and the Daniel I.C. Wang (1959) Faculty Research Innovation Fund at MIT. We thank the Mazumdar-Shaw International Oncology fellowship and the Koch Institute for supporting the research activities of H.N.

