## Supplementary material for "Accelerating Cell Culture Media Development Using Bayesian Optimization-Based Iterative Experimental Design": SI

### Supplementary Information

#### Retrospective Kernel Impact Evaluation

To retrospectively evaluate the impact of the kernel choice against the routine one hot encoding (OHE) approach, we used the data collected in this work and performed 50 random train–test splits of the data (80-20%). We trained the model on 80% data tested it on the remaining 20% and computed the root mean squared error in prediction (RMSEP), providing a distribution of 50 RMSEP.

$$\text{RMSEP} = \sqrt{\sum_{t=1}^{Ntest} \frac{\left(y_t^{obs} - y_t^{pred}\right)^2}{N}} \quad \text{(Eq. S1)}$$

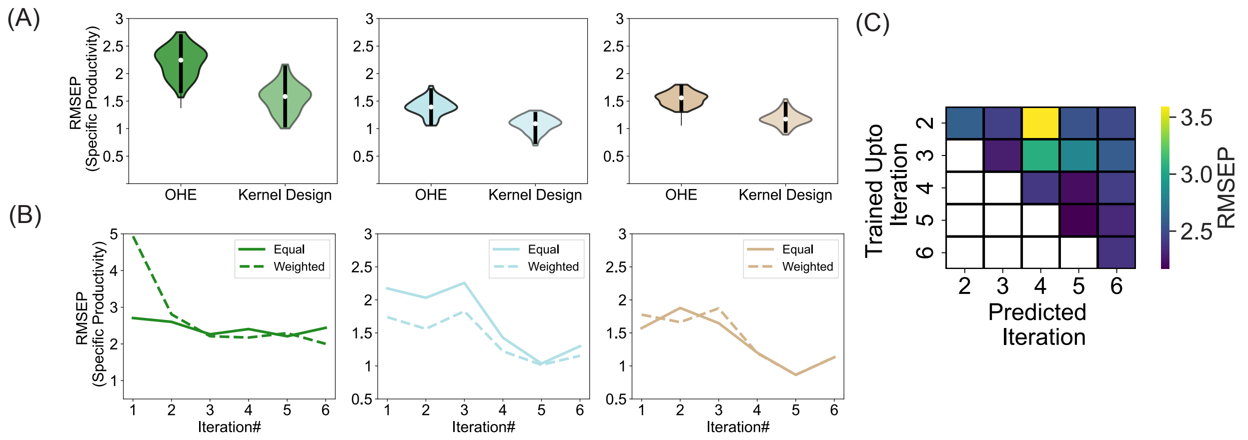


**Fig S1:** (A) Comparison of the RMSEP made by GP built using the OHE and designed kernel (B) Evolution of the predictive capability of the model in each iteration (C) Accuracy of the model in each iteration (x-axis) when accounting for progressively smaller subsets of data.

#### Converge Calculations

In this case, the model convergence was tracked through two approaches (i) the root mean squared error (RMSEP) and weighted RMSEP (ii) Progressive model prediction (Methods).

To check for convergence and the stopping criteria, we monitored the RMSEP and the weighted RMSEP for each iteration. In other words, during every iteration, we recorded the model prediction and compared it with the experimental observation obtained after performing the experiment. When the RMSEP made in two consecutive rounds is not significantly different it was determined as the indication of model convergence. To compute the weighted RMSEP the prediction with higher specific productivities ($SP_t$) were weighted higher by computing the weights as indicated in (Eq. S3).

$$RMSEP_{weighted} = \sqrt{\frac{\sum_{t=1}^{Ntest} w_{t*} \left(y_t^{obs} - y_t^{pred}\right)^2}{\sum_{t=1}^{Ntest} w_t}} \quad \text{(Eq. S2)}$$

$$w_t = \frac{SP_t}{\max(SP)} \quad \text{(Eq. S3)}$$

In addition, in every iteration, we also used a progressive model prediction approach. For instance, to predict the outcome of experiments in iteration 5, we would use models trained using data up to iterations 5, 4, and 3. We monitor the RMSEP made by each of these models trained on progressively smaller subsets of data. The point at which adding future iteration data is not crucial to the predictive performance was considered the checkpoint for model convergence.

The RMSEP ("Equal", Fig. S1 B) provides an estimate of the overall model accuracy, while the weighted RMSEP skews the metric to provide higher weightage to desired targets (higher specific productivities) in the calculation. As the iterations progress, both RMSEP and weighted RMSEP reduce indicating the improvement in the model accuracy with iterations 5 and 6 having comparable errors (Fig. S1 B). Also, from iteration 4 onwards there is an agreement between both metrics. Combined with the observation made about the design space (Fig. 5A), this can be attributed to the fact that the algorithm is predominantly exploiting the design space. With the progressive model prediction approach (Fig. S1 C), we also see that the RMSEP for a model trained with data up to iteration 4 performs comparably to a model trained with data up to iteration 6 thus providing an additional indication that the model is converged.

**Design space coverage of the extended design space considering the 9 factors**

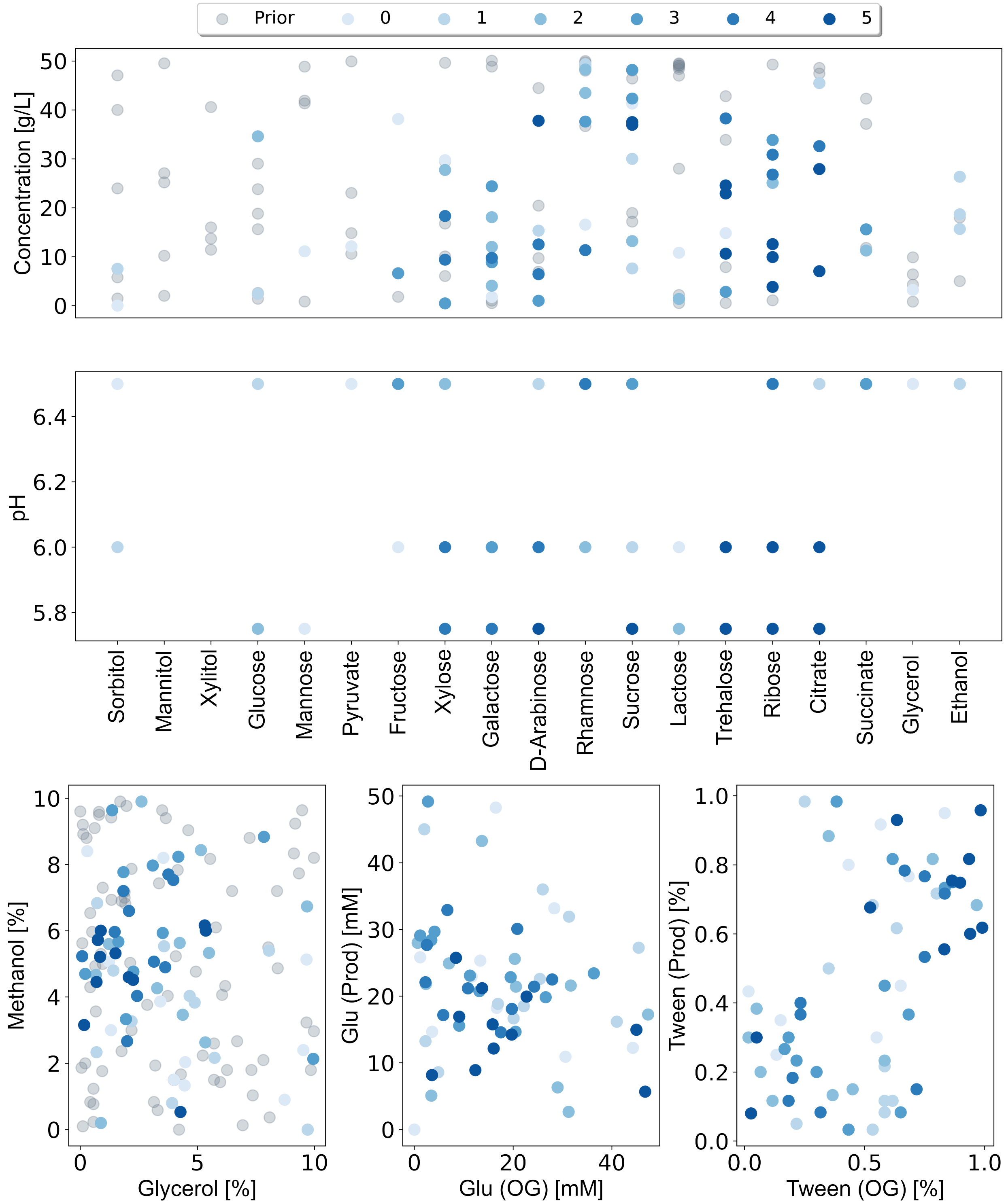


**Fig S2**: Evolution of the location of the experiments in the design space in the different iterations indicated through pairwise plots of the design factors.
